# Divergent DNA methylation signatures of juvenile seedlings grafts and adult apple trees

**DOI:** 10.1101/818690

**Authors:** Adrien Perrin, Nicolas Daccord, David Roquis, Jean-Marc Celton, Emilie Vergne, Etienne Bucher

**Affiliations:** IRHS (Institut de Recherche en Horticulture et Semences), UMR 1345, INRA, Agrocampus-Ouest, Université d’Angers, SFR 4207 QuaSaV, Beaucouzé F-49071, France; Plant Breeding and Genetic Resources, Agroscope, Nyon, Switzerland

**Keywords:** epigenetics, perennial plant, heritability, *Malus domestica*, sexual and asexual reproduction

## Abstract

Plants are continuously exposed to environmental perturbations. Outcrossing annual plants can adapt rapidly to these changes via sexual mating and DNA mutations. However, perennial and clonally reproducing plants may have developed particular mechanisms allowing them to adapt to these changes and transmit this information to their offspring. It has been proposed that the mechanisms allowing this plasticity of response could come in the form of epigenetic marks that would evolve throughout a plant’s lifetime and modulate gene expression. To study these mechanisms, we used apple (*Malus domestica*) as a model perennial and clonally propagated plant. First, we investigated the DNA methylation patterns of mature trees compared to juvenile seedlings. While we did not observe a drastic genome-wide change in DNA methylation levels, we found clear changes in DNA methylation patterns localized in regions enriched in genes involved in photosynthesis. Transcriptomic analysis showed that genes involved in this pathway were overexpressed in seedlings. Secondly, we compared global DNA methylation of a newly grafted plant to its mother tree to assess if acquired epigenomic marks were transmitted via grafting. We identified clear changes, albeit showing weaker DNA methylation differences. Our results show that a majority of DNA methylation patterns from the tree are transmitted to newly grafted plants albeit with specific local differences. Both the epigenomic and transcriptomic data indicate that grafted plants are at an intermediate phase between an adult tree and seedling and inherit part of the epigenomic history of their mother tree.

## Introduction

Epigenetic regulation of gene transcription is implemented by several covalent modifications occurring at the histone or DNA level without affecting the DNA sequence itself (Holliday and Pugh 1975). These modifications are termed epigenetic marks and can change throughout plant development. Some newly acquired epigenetic changes can also be inherited across generations (Hauser et al. 2011; Gutierrez-Marcos and Dickinson 2012; Kawashima and Berger 2014; Quadrana and Colot 2016). During their lifetime organisms may develop alternative phenotypes in response biotic and abiotic stresses (Madlung and Comai 2004; Mirouze and Paszkowski 2011; Köhler, Wolff, and Spillane 2012; Song, Irwin, and Dean 2013). These stimuli result in modifications in gene transcription which can be altered by epigenetic modifications (Manning et al. 2006; Schmitz et al. 2013; Kim and Zilberman 2014). Besides gene transcription changes, certain epigenetic marks have been shown to play key roles in DNA conformation and genome stability (Suzuki and Bird 2008; Hauser et al. 2011; Kim and Zilberman 2014). Indeed, DNA methylation has been shown to have a major role in transposable element (TE) silencing by reducing considerably the potential damage incurred by *de novo* TE insertions in the genome (Miura et al. 2001; Mirouze et al. 2009; Ito et al. 2011).

At the molecular level, DNA methylation consists in the covalent addition of a methyl group to cytosine nucleotide. In plants, DNA methylation occurs in three different cytosine contexts: CG, CHG and CHH (H= A, T or C) (Gruenbaum et al. 1981; Meyer, Niedenhof, and Ten Lohuis 1994; Finnegan et al. 1998; Chan, Henderson, and Jacobsen 2005). DNA methylation is established *de novo* or maintained by several DNA methyltransferase enzymes (Law and Jacobsen 2010), each having a specific role depending on the sequence context. In order to maintain DNA methylation following DNA replication that results in hemi-methylated DNA, the methyltransferases MET1 and CMT3 can copy DNA methylation patterns from the “ancestral” strand to the newly synthesized strand. This mechanism is called DNA methylation maintenance (Lindroth 2001; Schermelleh et al. 2007) and occurs at symmetric CG and CHG sequence contexts. However, for the CHH sequence context is no such template exists that may allow the DNA methylation maintenance mechanism. In this case, DNA methylation has to be restored by *de novo* methylation after each DNA replication cycle (Wassenegger et al. 1994; Chedin, Lieber, and Hsieh 2002). This pathway is called RNA-directed DNA methylation (RdDM) and requires small interfering RNAs (siRNA) (Herr et al. 2005; Kanno et al. 2005) to guide the DNA methylation machinery regions with sequence homology to the siRNAs.

From an epigenetic point of view, perennial plants are of particular interest as they have the potential to accumulate epigenetic modifications throughout their lifetime and may pass this information to the next generation. In addition, in the *Rosacea* family (Jung et al. 2019) numerous crops and ornamental plants are multiplied by asexual multiplication via grafting. This is interesting because in addition to the long lifetime of these plants, asexual multiplication involves only mitotic cell divisions (Verhoeven and Preite 2014) and thus presumably increases the chances of transmission of acquired epigenetic marks. If that was the case, epimutations could be quite common in grafted perennial plants. In contrast, during sexual reproduction meiosis can result in epigenetic reprogramming and therefore the loss of acquired epigenetic marks (Choi et al. 2002; Ibarra et al. 2012; Li, Kumar, and Qian 2018). In Arabidopsis, this reprogramming is the result of active DNA demethylation driven by DEMETER (DME) (Choi et al. 2002). Previous studies have suggested that this demethylation could contribute to the generation of totipotent cells (Slotkin et al. 2009; Gutierrez-Marcos and Dickinson 2012; Kawashima and Berger 2014) by alleviating gene silencing via active removal of DNA methylation. These modifications at the DNA methylation level are necessary for normal meiosis (Walker et al. 2018). The RdDM pathway remains active in the egg cell (Olmedo-Monfil et al. 2010). However, in the central cell of the mature female gametophyte and in the mature pollen sperm cell there is a decrease in RdDM activity (Kawashima and Berger 2014). This decrease releases the transcription of TEs, thus resulting in the production of siRNAs derived from those. These siRNAs have been reported to be transported into the egg cell (Han et al. 2000) to silence homologous loci in the maternal and paternal genomes (Han et al. 2000; Saze, Scheid, and Paszkowski 2003; Jablonka and Raz 2009; Feng, Jacobsen, and Reik 2010; Kawashima and Berger 2014). Based on these findings, one may assume that during sexual multiplication, meiosis would allow restauration of a specific DNA methylation level in these species, while during asexual multiplication mitosis would maintain epimutations.

In plants, inheritance of epigenetic marks has been widely investigated. Some studies point out the existence of broad epigenetic variations throughout wild populations of perennial and annual plants (Herrera, Medrano, and Bazaga 2016; Niederhuth et al. 2016; Wilschut et al. 2016). Other studies have demonstrated that epigenomic plasticity can allow environmental stress adaptation and improve response to future stresses (Herman and Sultan 2011; Herrera and Bazaga 2013; Medrano, Herrera, and Bazaga 2014; Colicchio et al. 2015). Finally, studies have suggested that epigenetic modifications induced by stress in a mother plant may improve stress response in their offspring (Agrawal, Strauss, and Stout 1999; Bilichak and Kovalchuk 2016; Ramírez-Carrasco, Martínez-Aguilar, and Alvarez-Venegas 2017). However, still little is known about heritable transmission of epigenetic marks in crops and more specifically in woody perennials like apple.

Apple (*Malus domestica*) is a major fruit crop in the world. In 2017, 130 million tons of fruit were produced on 12,3 million hectares (“FAOSTAT” 2017). In the Malus gender, tree multiplication for commercial orchards and conservation is performed via asexual multiplication. This vegetative multiplication (or clonal multiplication) obtained by grafting or budding ensures that all grafted trees originating from a particular cultivar are genetically similar. Scions of fruiting cultivars are grafted on rootstock to combine valuable agricultural traits. For instance, in addition to reducing tree size and modifying its architecture, grafting onto particular rootstocks is known to shorten the juvenile phase of the scion by promoting flower differentiation (Lane 1992). Scions can thus recover their ability to bloom 3 to 5 years after grafting (Lane 1992) while seedlings on their own roots may only start blooming after up to 8 years (Visser 1964). The juvenile phase is the first stage of development of new plants derived from sexual reproduction (Lavee et al. 1996). Juvenile phase length is highly variable among species, ranging from a few days, as in the *Rosa* genus (Hackett and Murray 2015) to more than 30 years in some woody plants (Rugini 1986; Bellini 1993; Meilan 1997). Certain phenotypic characteristics have been associated with the juvenile phase such as fast vegetative growth (Meilan 1997), low lignification of young shoots, short internodes, specific leaf shape (Lavee et al. 1996) and low trichome density. For instance, this phenotypic difference between juveniles and adults has previously been described in annual plants such as Arabidopsis (Telfer, Bollman, and Poethig 1997) or *Zea mays* (Poethig 2003), and perennials including the *Acacia* genus, *Eucalyptus globulus, Hedera helix, Quercus acutissima* (Wang et al. 2011) or in *Populus trichocarpa* (Critchfield 1960).

Here we investigated the transmission of epigenetic marks at the DNA methylation level using a recently completely sequenced apple doubled-haploid Golden Delicious line (GDDH13) (Lespinasse et al. 1999; Daccord et al. 2017). Taking advantage of this unique genetic material, we compared the effect of sexual and asexual multiplication at the phenotypic, gene transcription and DNA methylation levels. We present evidence that genome-wide DNA methylation levels are stable in apple independently of its multiplication mode. However, specific local variations in DNA methylation patterns involved in the regulation of key plant-specific gene regulatory networks such as photosynthesis were found and provide the basis for future studies on the role of epigenetics in tree aging.

## Results

### Phenotypic comparison of seedlings, young grafts and adult trees

We found that the GDDH13 doubled haploid apple showed a relatively high self-compatibility level as compared to the original ‘Golden Delicious’ variety from which it was derived. To prevent outcrossing and to produce self-fertilized GDDH13 seeds we covered trees with insect- and wind-proof cages during blooming time. Then we deployed bumblebees in the cages resulting in the production of hundreds of self-fertilized seeds. This unique material allowed us to study genetically identical seedlings and grafted plantlets derived from the very same parental tree. For that purpose, we simultaneously planted seedlings and grafted budwood from GDDH13 to ensure that the growing plants were of comparable size.

First, we studied the phenotypic differences between parental tree, grafts and seedlings on leaf samples in order to assess if the plants were in a juvenile or adult phase. Trichome density was the most noticeable phenotypic difference (Fig. 1). Leaves sampled from seedlings (Seedling) displayed a notably lower trichome density on their abaxial face (Fig. 1A) compared to the other samples. Leaves sampled from grafted plants (Graft) or from the original parental tree (Tree) showed a significantly higher trichome density (Fig. 1B-D).

**Figure 1:**
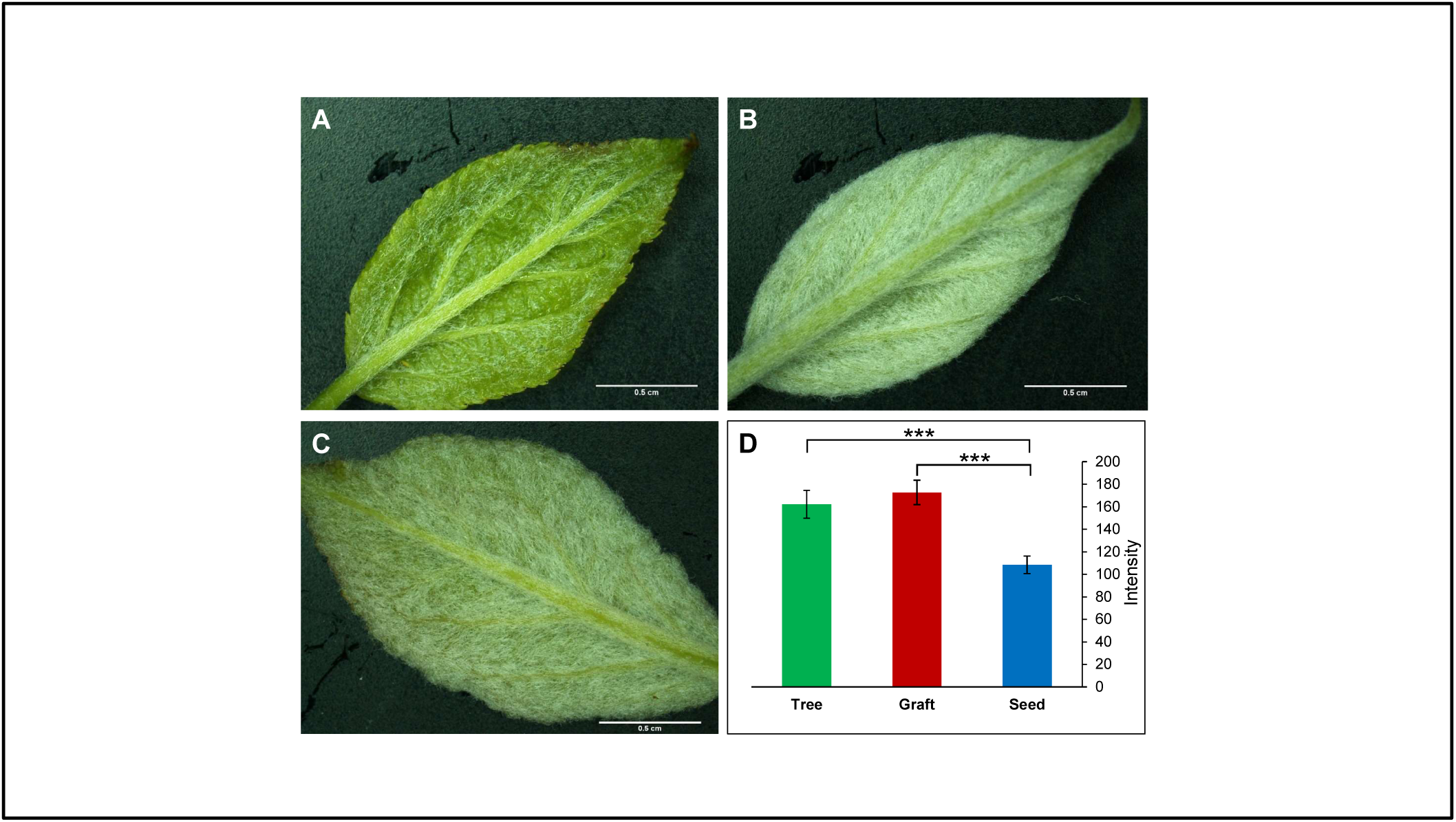
Leaf trichome density comparisons between seedlings, grafted plants and parental tree. Leaf pictures indicate visual differences in trichome density for seedlings (A), grafts (B) and donor tree (C). The graph in (D), represents results from light intensity measurements carried out on the abaxial face of leaves. High light intensity correlates with high trichome density. N = 60 (5 measures on 12 leaves) per sample. Statistical differences were evaluated by a Kruskall-Wallis test two by two. Asterix p-value: ***: 1‰.

In order to describe the gene regulatory mechanisms that may be underlying the observed phenotypic differences, we carried out transcriptomic analyses.

### Transcriptional profiles of seedlings, young grafts and adult trees

In order to identify genes related to the juvenile phenotype or genes displaying differential transcription levels in response to grafting, we performed a set of differential gene transcription analyses. We assessed steady state RNA levels by performing the following two comparisons: Tree versus Seedling (TvS) and Tree versus Graft (TvG). Transcriptomes were obtained using a custom-designed microarray that includes probes from all annotated GDDH13 genes and a fraction of TEs. We identified 6.943 and 7.353 differentially expressed transcripts (DETs) for TvS and TvG, respectively. Of these DETs, 5.695 were annotated as genes (DEGs) in TvS and 4.996 in TvG (Fig. 2A). In total these DEGs include 13,5% of all annotated gene on the microarray for TvS and 11,8% for TvG (Fig. 2A). For transcripts annotated as TEs, we identified 1.248 and 2.357 differentially expressed TEs (DETEs) in the TvS and TvG comparisons, respectively (Fig. 2B). These represent 5% of all annotated TEs on the microarray for TvS and 6,6% for TvG (Fig. 2B).

**Figure 2:**
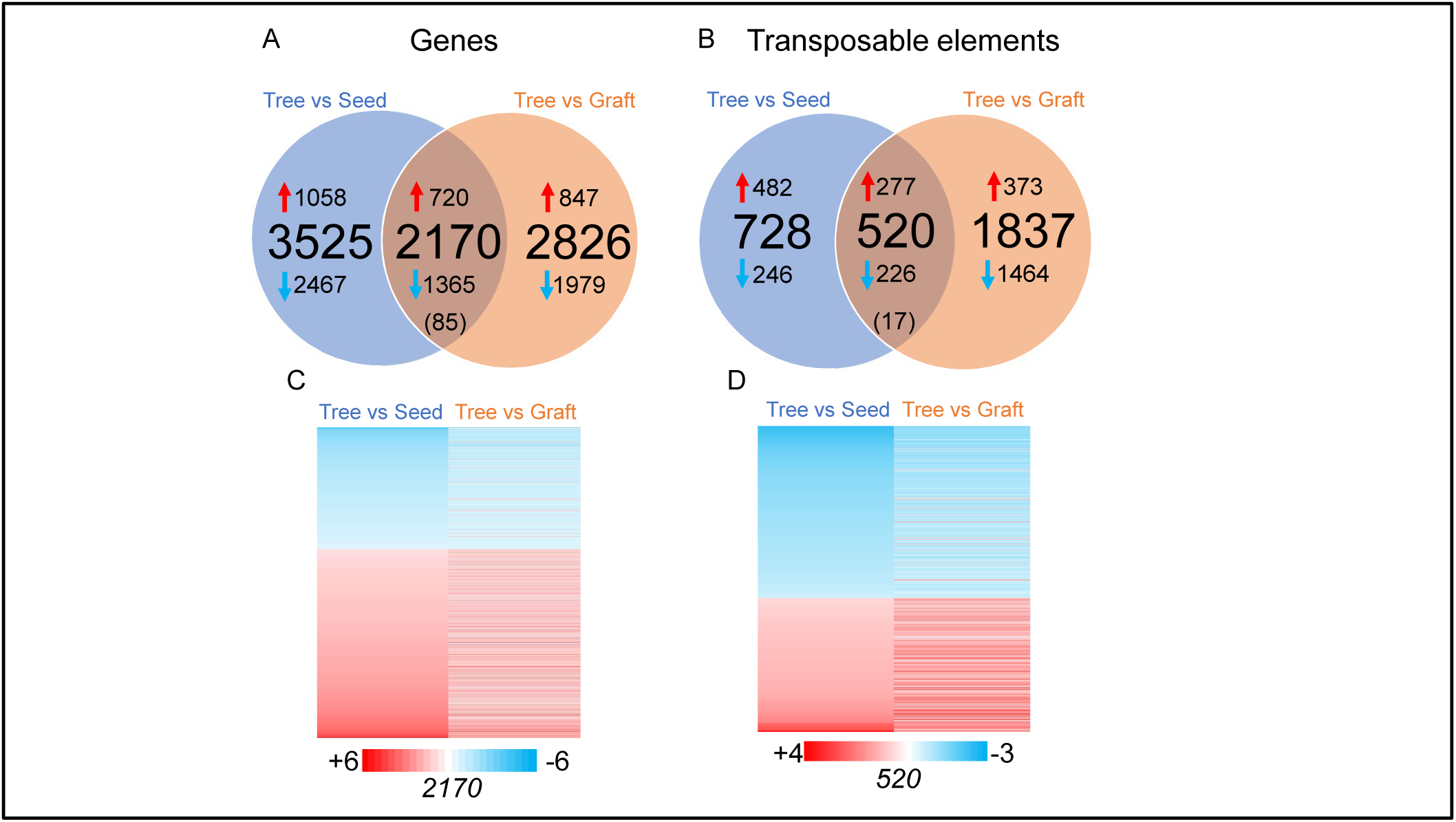
Transcriptome comparisons between seedling, grafts and donor tree. Graphical representation of the number of differentially expressed transcripts in the different comparisons. (A) Venn diagram showing differentially transcribed genes (DEGs) in the comparisons TvS and TvG. (B) Venn diagram depicting differentially expressed TEs (DETEs) in the comparisons TvS and TvG. The central number in brackets represent common DETs displaying alternative pattern of transcriptional regulation. In (C) and (D) the heat maps depict transcription ratios of common DEGs (C) and DETEs (D). Numbers of DETs in each heat map are indicated below it. Fold change ratios are shown in the color scale bar.

Overall, DEGs displayed a tendency towards down regulation in Tree compared to Seedling and Graft (Fig. 2A). However, for TEs only the TvG comparison followed the same pattern, since up- and down-regulated TEs were more equally distributed in the common DETEs group. DETEs specific to TvS displayed a tendency to be up regulated in Tree.

Focusing on the common DEGs between TvS and TvG, we observed two groups (Fig. 2A and C). The first group is composed of the 2.085 DEGs displaying a similar regulation pattern: 1.365 and 720 DEGs were down and up regulated in TvS and TvG, respectively. In the second smaller group, only 85 DEGs displayed an opposite trend: these transcripts were down regulated in Tree in TvS, but up regulated in Tree in TvG. Similarly, we observed two groups for DETEs (Fig. 2B and D). 277 DETEs were up regulated in Tree in both TvS and TvG, and 225 DETEs were down regulated in Tree in both comparisons. Only 17 DETEs displayed an opposite transcript accumulation patterns compared to the general trend.

To study the main gene regulatory pathways represented in the differential transcription data we used the GDDH13 gene annotation of *Malus domestica* (v1.1) combined with the MapMan software (Lohse *et al*., 2014; Fig. 3A). We also considered the TE class repartition as previously described in Daccord *et al*., (2017) (Fig. 3B). We observed variations in class size between TvS and TvG. The most notable variations size were observed for: photosynthesis (9% of variation in total DEGs in TvS and only 1% in TvG), cell cycle (2% in TvS and 9% in TvG), solute transport (9% in TvS and 4% in TvG), cytoskeleton (1% in TvS and 6% in TvG), RNA biosynthesis (13% in TvS and 18% in TvG), RNA processing (2% in TvS and 6% in TvG) and chromatin organization (2% in TvS and 5% in TvG).

**Figure 3:**
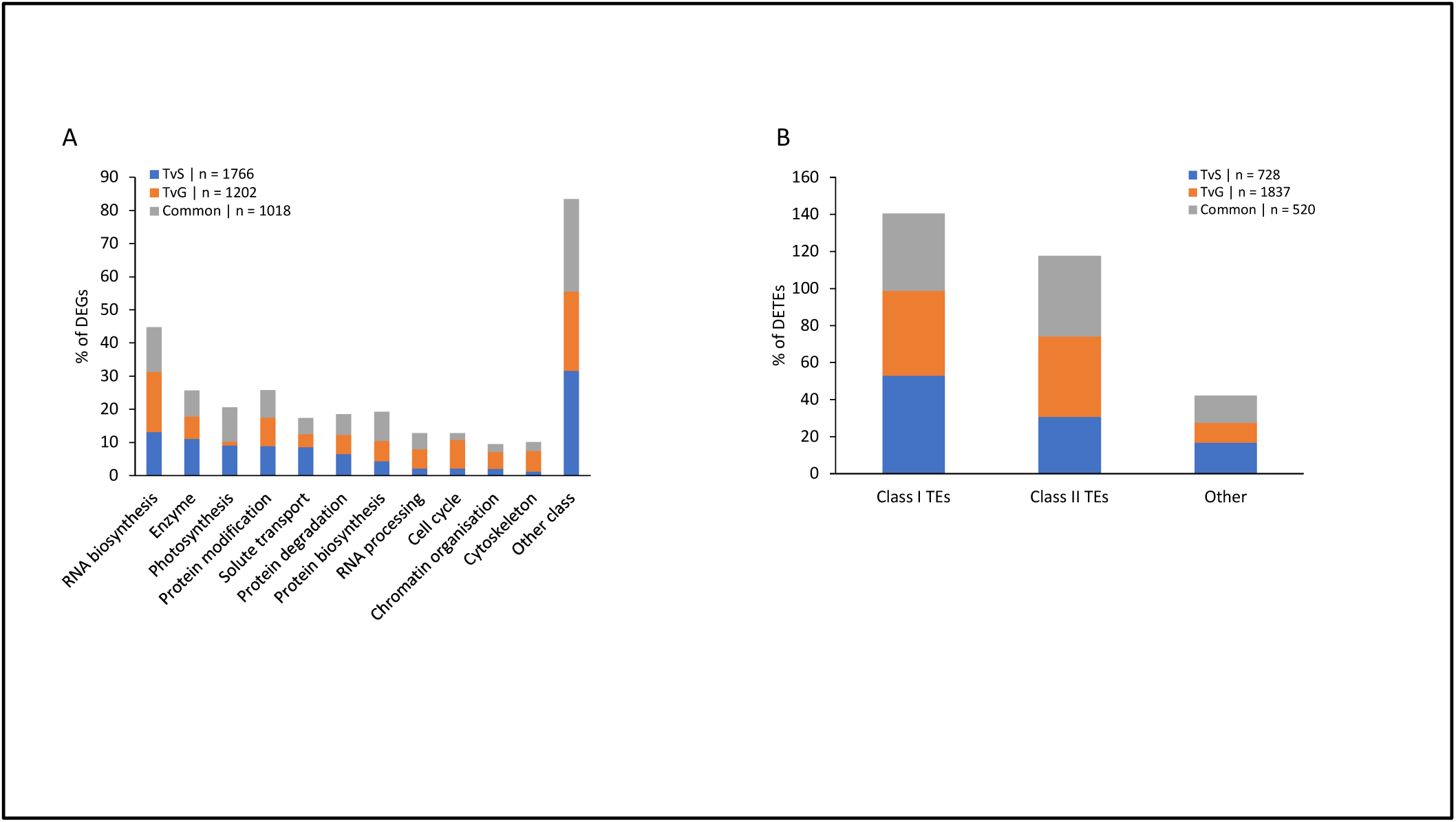
Classification of differentially expressed transcripts. (A) percentage of DEGs in each comparison in function of the gene classification according to Lohse et al. (2014). (B) percentage of DETEs in each comparison in function of TEs classification according to Daccord et al. (2018). Classes represented by less than 5% in the three condition were summed up in “Other class”.

In order to identify overrepresented classes of genes that could be linked to either the adult or the juvenile phase, we performed an enrichment analysis with MapMan using our DEGs as input data (Supplemental Tab. S1). In the TvS comparison, seven functional categories were overrepresented including coenzyme metabolism, terpenoids metabolism, chromatin organization, squamosa binding protein (SBP) family transcription factor, protein biosynthesis, peptide tagging in protein degradation and enzyme classification. Eleven classes are overrepresented in the TvG comparison (Supplemental Tab. S1), including secondary metabolism, chromatin organization, cell cycle, RNA processing, protein biosynthesis, peptide tagging, cytoskeleton, cell wall, solute transport, and enzyme classification.

**Table 1:** DMR distributions according to context and methylation changes. Number and percentage of hyper- and hypomethylated DMR in Tree sample in each comparison (TvS and TvG).

| Context | Tree vs Seed |  |  | Tree vs Graft |  |  |
| --- | --- | --- | --- | --- | --- | --- |
|  | Hypermethylated | Hypomethylated | Σ | Hypermethylated | Hypomethylated | Σ |
|  | Number (%) | Number (%) | Number (%) | Number (%) | Number (%) | Number (%) |
| CHH | 203671<br>(90,06) | 22482<br>(9,94) | 226153<br>(98,74) | 92820<br>(63,01) | 54485<br>(36,99) | 147305<br>(95,42) |
| CHG | 175<br>(17,00) | 854<br>(83,00) | 1029<br>(0,45) | 209<br>(8,20) | 2339<br>(91,80) | 2548<br>(1,65) |
| CG | 411<br>(22,20) | 1440<br>(77,80) | 1851<br>(0,81) | 480<br>(10,63) | 4037<br>(89,37) | 4517<br>(2,93) |
| Σ | 204257<br>(89,18) | 24776<br>(10,82) | 229033<br>(100) | 93509<br>(60,57) | 60861<br>(39,43) | 154370<br>(100) |

Next, we considered the TE class repartition in our DETE list (Fig. 3B). We did not find large variations in class repartition among the comparisons. Class I TE represented 53% of DETEs on the microarray in TvS and 46% in TvG. Concerning class II TEs we found 31% and 43% of DETEs in TvS and in TvG respectively.

Altogether, our analyses show that the two sexual and asexual tree propagation methods investigated here had a significant effect on gene and TE transcription in GDDH13.

### Global DNA methylation analysis of seedlings, young grafts and adult trees

To investigate how DNA methylation marks are transmitted through mitosis as compared to meiosis, we assessed the DNA methylation levels in Seedling, Graft and Tree samples at the genome-wide level by using whole genome bisulfite sequencing (WGBS). First, we compared the genome-wide DNA methylation levels at cytosines in the three sequence contexts (CG, CHG, CHH). Our primary investigation indicated that there was no significant difference in cytosine methylations averages, in any of the contexts, among the tested samples (Fig. 4A). Next, we computed and identified differentially methylated regions (DMR) between Seedling, Graft and Tree. Overall, we identified 229.033 DMRs in TvS and 154.370 in TvG (Fig 4B). We also investigated DMRs close to genes (Gene-DMRs) or TEs (TE-DMRs). These DMRs are defined by their relative proximity to genes or TEs. For this purpose, we selected DMRs located within 2.000 bp in 3’ or 5’ of annotated genes or TEs. We identified 48.651 and 18.789 Gene-DMRs in TvS and TvG, respectively. For TE, we identified 124.025 and 97.330 TE-DMRs in TvS and TvG, respectively (Fig. 4B).

**Figure 4:**
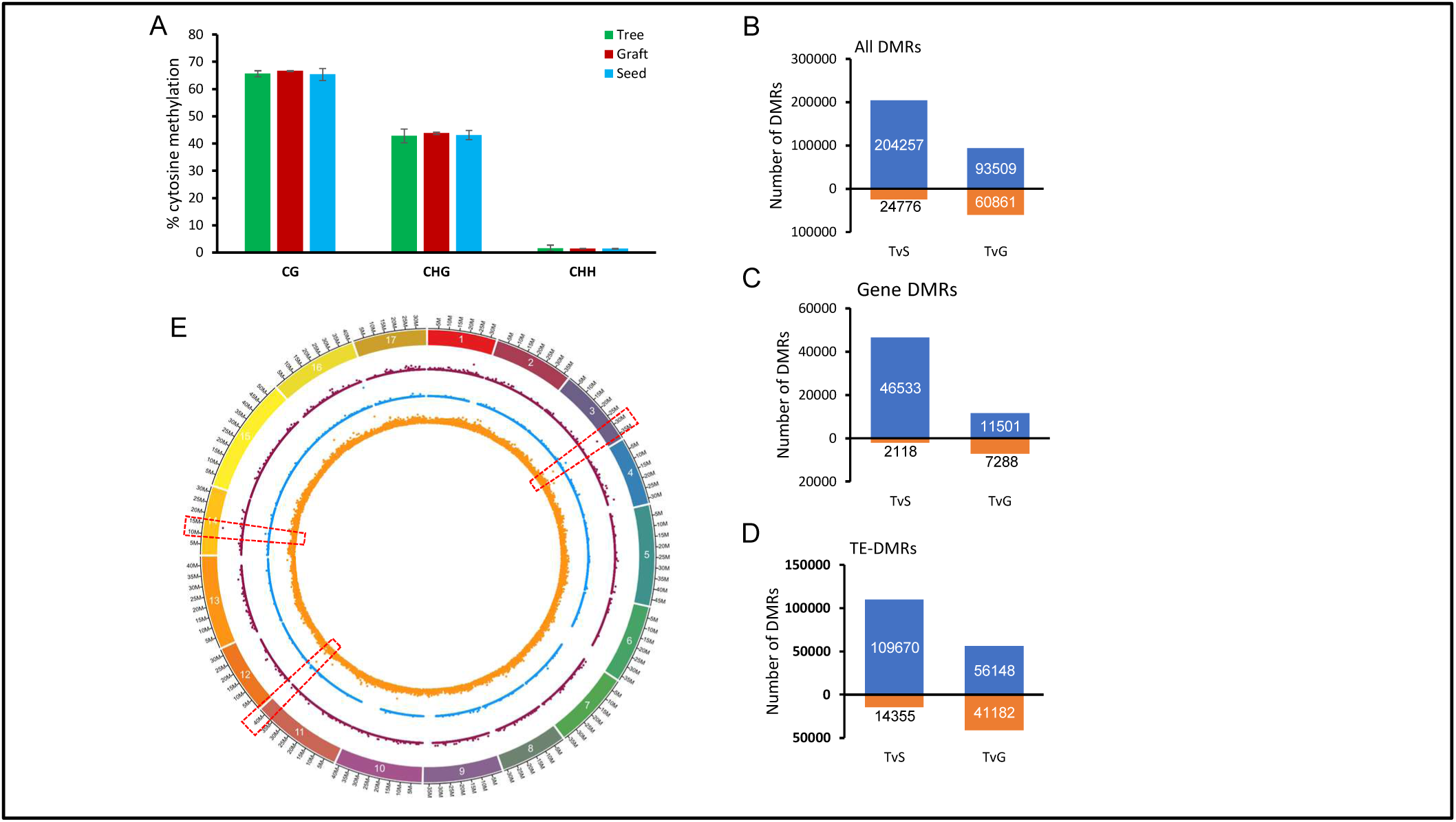
Global overview of DNA methylation differences between seedlings, grafts and trees. (A) Histogram presenting the genome wide cytosine methylation level (in percentage) of the three methylations context (CG, CHG and CHH). Student test was performed to evaluate differences and the results (B, C and D). Histograms representing the number of DMRs for each comparison: hypermethylated (above 0, in blue) or hypomethylated (below 0, in orange) in the Tree samples for all DMRs (B), Gene-DMRs (C) and TE-DMRs (D). DMRs in all sequence contexts were counted and values are indicated in graph. (E) density plot of number of DMRs in 50 kb windows on the GDDH13 genome for TvS (see supplemental figure S1 for TvG). In red, DMRs in the CG context, in blue for the CHG context and in orange the CHH context. Each point represent the number of DMRs in a 50kb window of the genome. Red dashed boxes indicate the presence of DMR hot spots.

We found that in each comparison, in genes, TEs or other genomic loci, DMRs were largely hypermethylated in Tree (Fig. 4B). Indeed 89% and 61% of DMRs in the three contexts were hypermethylated in TvS and in TvG respectively. Moreover, a vast majority of DMRs were identified in the CHH context (95% and 99% in TvG and TvS, respectively; Tab. 1). Overall, DMRs tended to be hypermethylated in Tree in the CHH context (90% in TvS and 63% in TvG) and hypomethylated in Tree in the CG and CHG contexts (93% in TvS and 92% in TvG) (Tab. 1). To identify whether DMRs were equally distributed along the genome, or were regrouped within hot spots, we computed the DMR density for the individual contexts as shown in Fig. 4C. Overall, we found that DMRs to be equally distributed all along the apple chromosomes, with some regions displaying a higher enrichment (Fig. 4C, red boxes).

In order to quantify and compare DNA methylation levels we compared DNA methylation changes (δmC) within DMRs in each sequence context (Fig. 5). Overall, we identified significant differences in δmC for the CHG and CHH and not for the CG sequence contexts. Interestingly, in the CHG context, the δmC value was higher in TvG (9.8%) than in TvS (5.4%) for hypermethylated DMRs in Tree. For hypomethylated DMRs in Tree, the δmC value was higher in TvS (9.6%) than in TvG (8.1%). In the CHH context, we observed that the δmC value was higher in TvS (5.8%) than in TvG (4.9%) for hypermethylated DMRs in Tree, and lower in TvS (3%) than in TvG (5.8%) for hypomethylated DMRs in Tree. From these results, we conclude that the transmission of cytosine methylation from Tree to Seed is different to the one from Tree to Graft depending on the cytosine sequence context.

**Figure 5:**
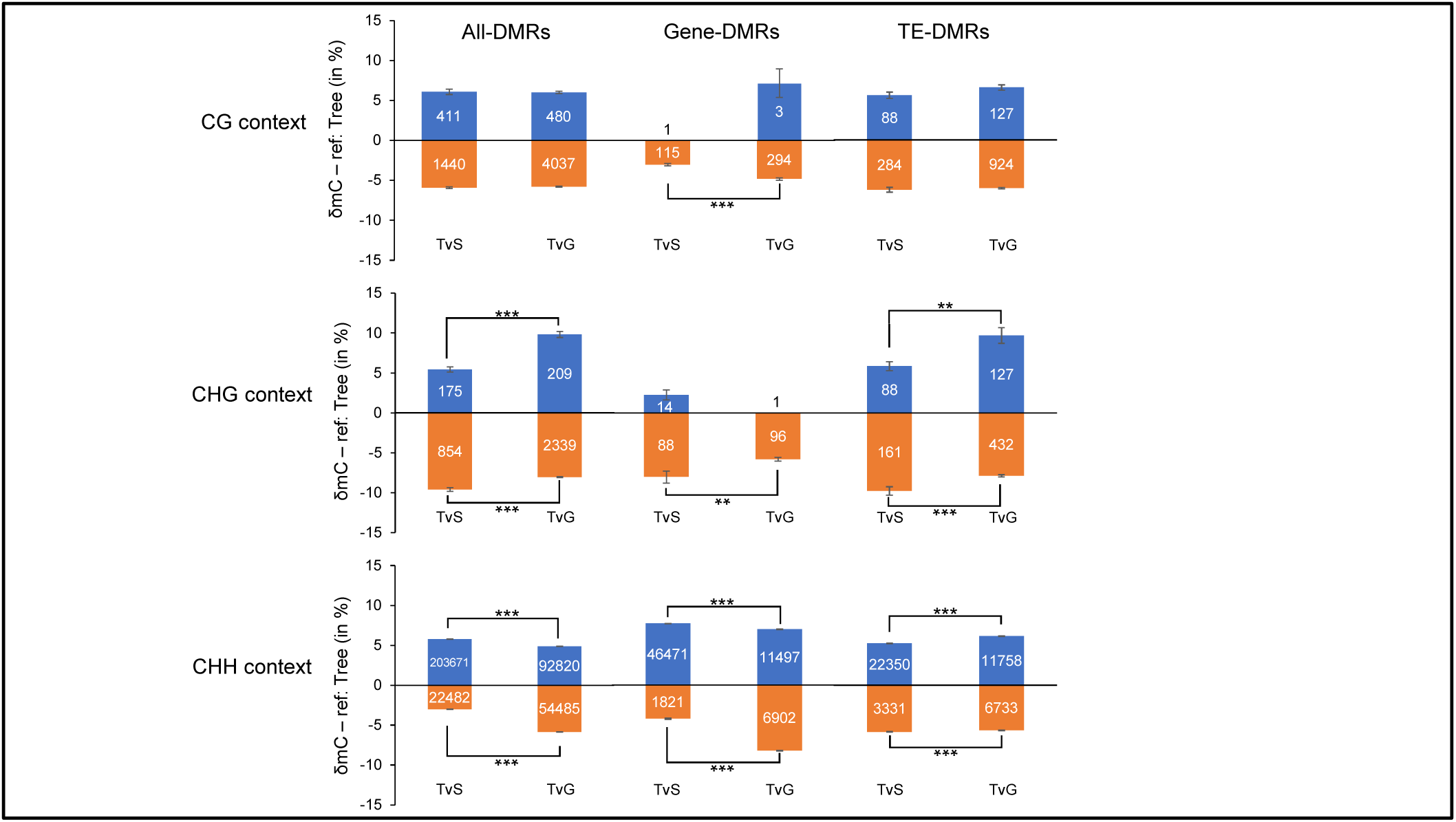
Levels of DNA methylation changes in gene and TE annotations. Histograms depicting DMR methylation variations (δmC) between samples separated by sequence context and functional annotation. All DMRs are presented in the All-DMRs column, genes and TEs in the Gene-DMRs and TE-DMRs, respectively. DMRs were filtered by p-value and SDA (standard deviation average) in accordance to a fixed threshold (Table S2). Student test was performed to evaluate differences in δmC, results are represented by an asterix depending on the p-value threshold: *: 5%; **: 1%; ***: 1‰. δmC: delta of methylation. The Tree sample was taken as reference to define the hyper- or hypomethylated state of DMRs.

For DMRs located in genic regions (Gene-DMRs, Fig. 5) we observed that there were less DMRs in the CG-CHG (359 for TvS and 390 for TvG) contexts than in the CHH context (48.292 for TvS and 18.399 for TvG). Gene-DMRs in CG and CHG context were almost all hypomethylated in Tree in both comparisons. Indeed, 99% of Gene-DMRs in the CG context were hypomethylated in both comparisons. 86% and 99% of Gene-DMRs were hypomethylated in CHG in TvS and TvG, respectively. This is consistent with the observations we made for the All-DMRs group (Tab. 1). Conversely, 96% and 62% of Gene-DMRs in the CHH context were hypermethylated in Tree for TvS and TvG, respectively.

While studying the DNA methylation changes, we found that in the CG and CHH contexts, the δmC values of hypomethylated Gene-DMRs were smaller in TvS (3.0 and 4.2% respectively) than in TvG (4.8 and 8.2% respectively). However, for hypomethylated Gene-DMRs in the CHG and CHH contexts in Tree the δmC value was higher in TvS (8.0 and 7.7% respectively) than in TvG (5.8 and 8.2% respectively) following the overall trend observed for All-DMRs. These observations indicate towards a contrasted sequence context specific pattern of DNA methylation differences.

For DMRs located in TE annotations (TE-DMRs, Fig. 5), our observations were similar to the results for Gene-DMRs. Overall most TE-DMRs were hypomethylated in Tree in the CG (76% for TvS and 88% for TvG) and CHG (65% for TvS and 77% for TvG) contexts, and hypermethylated in the CHH (87% for TvS and 64% for TvG) context. We did not find significant differences in δmC values for the CG context. For TEs, the δmC value of hypermethylated TE-DMRs was smaller in TvS (6.2 and 5.6% respectively) than in TvG (10.3 and 6.5% respectively) and higher for hypomethylated TE-DMRs in TvS (10.4 and 6.2% respectively) as opposed to TvG (8.4 and 6.0% respectively).

Even though there were no strong global differences in DNA methylation level between the samples analyzed here, we found significant local differences. The majority of DMRs were in the CHH context with a tendency to be hypermethylated in Tree.

### Classes of genes enriched with DMRs

To identify genes belonging to particular functional categories and presenting DMRs in their proximity, we used the aforementioned GDDH13 annotation in MapMan and the TE annotation as previously used in our transcriptomic analysis. Here we only considered Gene-DMRs and TE-DMRs in the CHH context. We excluded DMRs associated with the CG and CHG context here analysis due of their very limited number (Supplemental Tables S2 and S3). For the following, we termed as DEG-DMRs genes that we found to be differentially transcribed and containing or being close to DMRs. Similarly, TEs identified as DETEs and being associated with TE-DMRs were termed DETE-DMRs.

As expected, we found the seven classes that we previously identified in DEGs analysis: RNA biosynthesis, protein modification, enzyme family, protein degradation, solute transport, photosynthesis and protein biosynthesis (Fig. 6A). We did not find differences in the proportion of gene classes between TvS and TvG.

**Figure 6:**
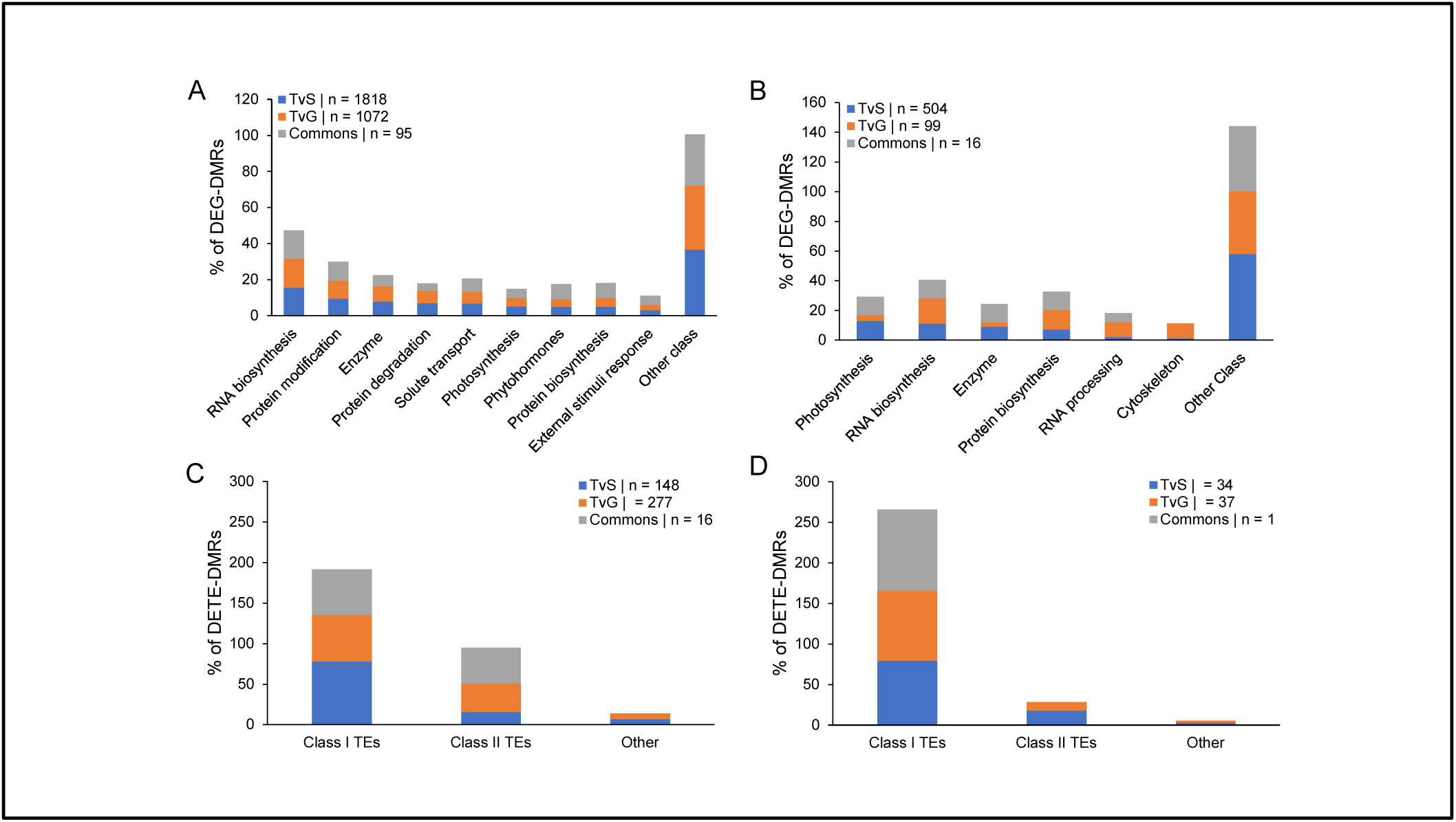
Classification of differentially expressed genes that are associated to DMRs. Histograms describing the percentage of DEG-DMRs (A and B) and DETE-DMRs (C and D) in the respective comparisons in function of gene or TEs classification. Only DMRs in the CHH context are presented here. In (A) and (C) all DEGs- and DETEs-DMRs were used while in (B) and (D) we only considered DEGs and DETEs with differential transcription ratio greater than 1.5 in absolute value. Gene classes representing less than 5% (A) or 10% (B) of the total in the three conditions were summed up in “other class”.

For DETE-DMRs (Fig. 6C) we observed a smaller proportion of Class I TEs in TvG (57,4%) compared to TvS (77,7%), while for class II TEs we found 35.4% for TvG and 15.5% for TvS.

### Relationship between DNA methylation and transcription

Next, we associated Gene- and TE-DMRs to our microarray transcriptome data and the aforementioned gene classes are defined according to the Mapman annotation of genes and to the TE annotation previously used to analyze DEGs and DETEs. For this analysis we applied a threshold and kept only transcripts with differential expression ratios above 1.5 and below −1.5 in order to better identified pathways or genes to work with.

We found 520 DEG-DMRs in TvS and 115 DEG-DMRs in TvG (Fig. 6C), 35 DETE-DMRs in TvS and 38 DETE-DMRs in TvG (Fig. 6D). We investigated genes and TEs classes’ repartition for DEG- and DETE-DMRs. Of the eleven classes found in DEGs analysis (Fig. 3A), here, we found only six gene classes representing only slightly more than 5% of all DEG-DMRs. These including the classes photosynthesis, RNA biosynthesis, enzyme family, protein biosynthesis, RNA processing and cytoskeleton (Fig.6B).

We did not observe notable shifts within the classes’ repartition between TvS an TvG for DETE-DMRs (Fig. 6D).

Finally, we investigated the link between DMRs and DEGs. As previously we only considered DEG-DMR in the CHH context because of the low number of DEG-DMR we found in the CG and CHG context. We noticed that in both comparisons the majority of DMRs in DEG-DMRs were located in gene promoters (539 for TvS and 114 for TvG), followed by terminator region (284 for TvS and 58 for TvG) finally followed by those present in gene bodies (139 for TvS and 42 for TvG) (Fig. 7A, 7B. See examples of these DEG-DMRs in Supplemental fig. S2).

**Figure 7:**
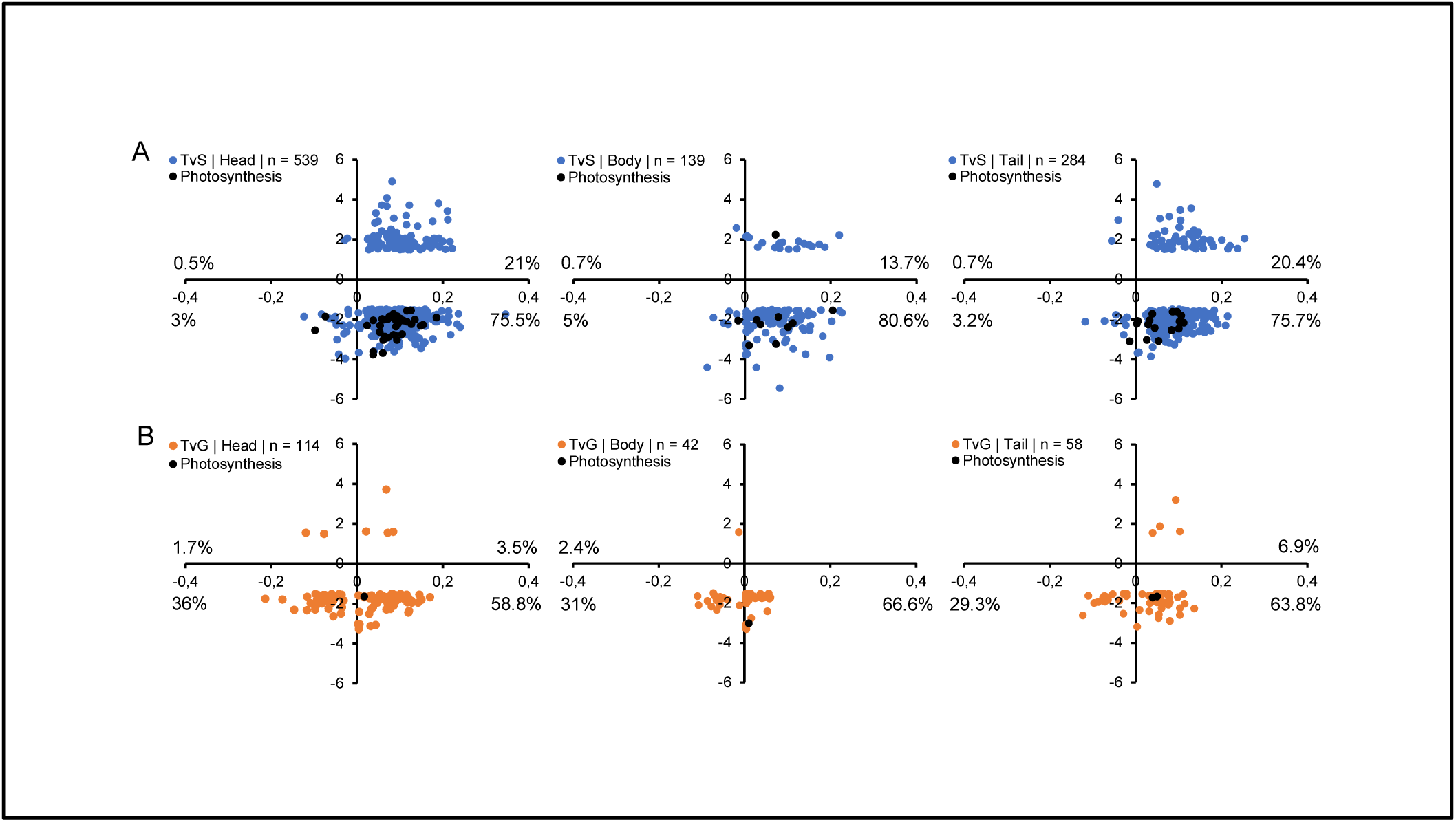
Relationship between transcription ratio and DNA methylation variation. Scatterplot representing DEG-DMRs in TvS (A) and TvG (B) in the CHH context, X axis represents δmC and Y axis represents gene expression ratios. In blue/orange are shown all DEG-DMRs and in black the ones specifically associated to photosynthesis. Numbers of DEG-DMRs used in each graph are indicated in the legend and percentages indicate number of DEG-DMRs in each corner of the graph. We separated DEG-DMRs in function of the position of the DMRs related to the corresponding gene (head = promoter, body, tail = terminator). Here we included the unannotated gene class “35” not present in Fig. 6.

Our data also indicate that, independently of the DMR position relative to a gene, hypermethylated DMRs were associated with a gene down-transcription in the Tree sample (Fig. 7A & B).

Among the classes of differentially transcribed genes associated with DMRs, we observed that genes associated with photosynthesis were mostly both hypermethylated and downregulated in the Tree sample. Indeed 92% of Gene-DMRs associated to photosynthesis pathway present this pattern in TvS and 100% of them in TvG.

Our results indicate that in the CHH context, hypermethylation of a DNA sequence in the proximity of a gene reduce the level of transcription of that particular gene.

## Discussion

### Newly grafted plants are at an intermediate state between adult tree and juvenile seedling

Phenotypic differences between juvenile and adult plants are commonly observed at the leaf level (Lavee et al. 1996). In our study we observed that leaves of seedlings displayed a low trichome density compared to grafted plants and to the donor tree (Fig. 1). As previously reported by others, this phenotype can be associated to the juvenile phase (Basheer-Salimia 2007) and the grafted plant seems thus closer to the adult tree than to the juvenile seedling from that point of view. Nonetheless, newly grafted plants show a contrasted ability to flower. In the grafting process, a mature bud (able to flower or quiescent) is placed on a short-rooted stem (rootstock). The number of nodes between the apical bud and the rootstock is drastically reduced to 1 or 2 nodes. After their first year of growth, buds are in a mature adult state but are unable to initiate flowers and to bear fruits because of an insufficient number of nodes (less than 77) in the stem, a limit previously described as a transition phase between juvenile and adult apple tree (Zimmerman 1973; Hanke et al. 2007; X. Z. Zhang et al. 2007).Thus, grafted plants are not adult plants from a physiological point of view.

Here, we wanted to study the molecular changes that occur during propagation via grafting and by seed formation. First, we compared the transcription profiles in three different stages: seedlings, grafted plants and the donor tree, taking advantage of our genetically identical material growing under highly similar conditions. Globally we observed a lower transcription level for the majority of the DEGs and DETEs in the adult tree compared to grafts or seedlings. This correlates well with the previously reported decrease in gene transcription in mature plants, compared to juvenile plants (Murray, Smith, and Hackett 1994; Hand et al. 1996; Ryan, Binkley, and Fownes 1997). Furthermore, the common DEGs identified in Tree versus Seedling and Tree versus Graft comparisons as repressed could be correlated to high vegetative growth in younger stages such as seedlings and grafted plants, and thus be transcribed at a lower level in mature apple tree, as observed in Day, Greenwood and Diaz-Sala, (2002) (Day, Greenwood, and Diaz-Sala 2002). Thus, the transcriptome of newly grafted plants showed similarities with the one obtained from seedlings but also with adult trees. These observations are in line with previous studies on other woody plant (Murray, Smith, and Hackett 1994; Hand et al. 1996; Ryan, Binkley, and Fownes 1997; Day, Greenwood, and Diaz-Sala 2002).

Overall, our findings indicate that young grafted plants are at the interface between a juvenile seedling and an adult mature tree (Zimmerman 1973; Hanke et al. 2007) from a morphological and transcriptomic perspective.

This intermediate condition of newly grafted plants is confirmed from a physiological point of view. Indeed, we identified differences in gene class repartition of DEGs between TvS and TvG (Fig. 3A) which included classes photosynthesis, RNA processing, chromatin organization and cell cycle. And concerning genes related to photosynthesis we found that they represented 9% of DEGs in TvS but only 1% in TvG. This is consistent with the fact that the photosynthetic pathway has previously been described as differentially regulated between juvenile and mature reproductive plant, especially in woody plants (reviewed in Bond, 2000), and is known as a physiological process subjected to many modifications from juvenile to mature phase (Greenwood 1995). As juvenile, seedlings undergo broader transcriptomic changes compare to grafts and trees. This can be associated to an age-related gene transcription pattern previously described for photosynthesis related genes in other woody plant such as *Pinus taeda* (Greenwood 1984), *Larix laricina* (Hutchison et al. 1990), *Picea rubens* (Rebbeck, Jensen, and Greenwood 1993) and in *Quercus* gender (McGowran, Douglas, and Parkinson 1998).

This intermediate condition of newly grafted plants between juvenile seedlings and adult tree was also observed at specific loci at the DNA methylation level in the CHH context. Indeed, overall a hypermethylation of the CHH-DMRs was observed in trees compared to grafts, which was less extended (62% in TvS) compared to the nearly total hypermethylation of CHH-DMRs observed in the Tree sample compare to Seedlings (96% in TvG).

### DMRs influence neighboring gene transcription

Previous reports established a correlation between DNA methylation and the repression of gene transcription, particularly in the model plant Arabidopsis (X. Zhang et al. 2006; Zilberman et al. 2007). In this study, we investigated a possible link between DNA methylation and gene transcription changes in *M. domestica*. For that purpose, we associated DMRs with their neighboring DEG in order to investigate the effect of methylation on gene transcription. We found that in the CHH context, genes with closely located hypermethylated DMRs (in Tree sample) often displayed a lower gene transcription level in Trees compared to Seedlings or Grafts (Fig. 7). This was particularly the case for photosynthesis related genes (Fig. 7). Our data indicate thus that cytosine methylation in the CHH context seems to be involved in regulating the transcription of these genes.

We did not only observe local changes in DNA methylation, but also contrasted levels of DNA methylation changes (δmC). Indeed, we found significant differences at the δmC level between both comparisons, particularly in the CHG and CHH contexts (Fig. 5). For methylation in the CHH context, we observed that, even if the difference in δmC was significant between TvS and TvG, it is not very high between the comparisons but also within comparisons. Indeed, the highest δmC was on average above 8% for hypomethylated Gene-DMRs in TvG. But we also found that this relatively small methylation variation was enough to find relationship with gene transcription changes (Fig. 7).

## Conclusion

In this study we compared the transmission of epigenetic marks and their potential effects on transcription during sexual and asexual reproduction in apple.

First, we identified a phenotypic change (Fig. 8) that was associated with adult plant phase and confirmed that grafting is not comparable to a complete rejuvenation process, as observed in seedlings. In our transcriptomic analysis we showed gene level transcription differences of the tree compared to seedlings and grafts (Fig. 8). In particular, we found that the transcription level of genes related to photosynthesis was relatively high in seedlings compared to the tree, while newly grafted plants displaying an intermediate transcription level (Fig. 8).

**Figure 8:**
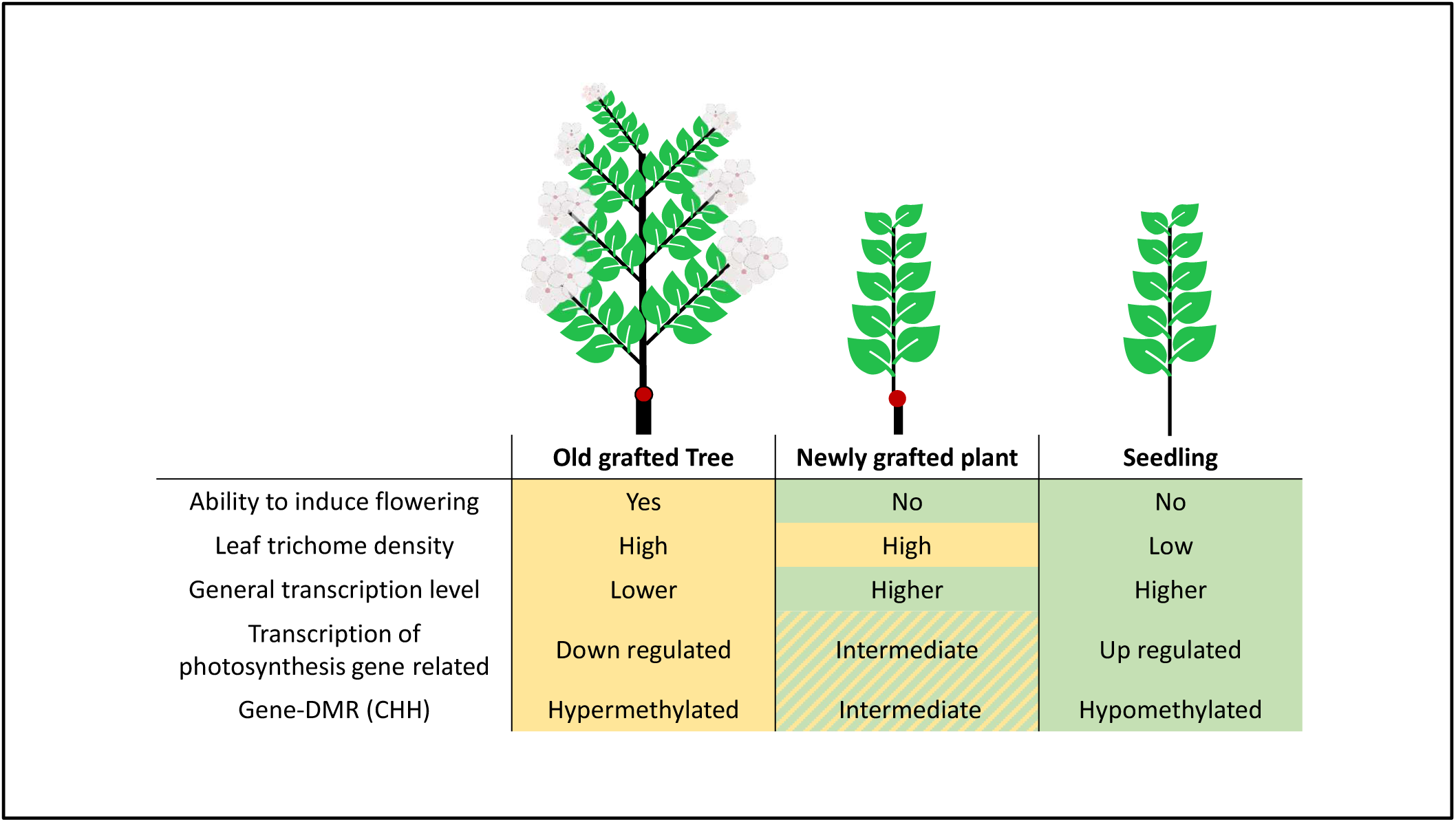
General overview of the main results of this study concerning physiological and molecular changes occurring during sexual and an asexual multiplication. The red dot represents the grafting point between scion and rootstock (larger line weight). Shared aspect between plants are highlighted by background colours.

Analysis of the methylation data indicated that at the genome scale, the level of methylation in all three samples was similar. However, we were able to identify DMRs particularly in the CHH context. This result indicates that methylation reprogramming during meiosis may not affect the global methylation level of the genome, but rather modify particular regions of the genome, presumably allowing the seedling to increase its competitiveness. This observation was particularly striking regarding genes associated with photosynthesis. As found in transcriptomic analysis, the methylome data indicated that grafted plants were at the interphase between the tree and the seedlings.

Globally, our results indicate that, from a physiological, transcriptomic and epigenomic standpoint, newly grafted plants are at the interphase between a tree and a seedling, displaying characteristics that are particular to both the mature and the young immature stages of the plant.

## Materials and Methods

### Plant material

*Malus domestica* materials were obtained from ‘GDDH13’ (Lespinasse et al. 1999) line (X9273). Grafted plant (called “Graft” in this paper) materials were obtained by grafting budwood of ‘GDDH13’ orchard tree (2001) on the rootstock ‘MM106’. Seedling materials (called “Seedling” in this paper) were obtained by self-fertilization of ‘GDDH13’ tree in 2017. Seed dormancy was removed by 3 months of cold stratification before sowing. Homozygous state of seedling was confirmed by PCR, using SSR markers on the seedling samples used in this work. A clone of the original ‘GDDH13’ from orchard, grafted onto an MM106 rootstock in 2007 and placed in the greenhouse in 2016 was used as reference mature adult tree (called “Tree” in this paper).

### Phenotyping

Nine young leaves were harvested for each sample and time point. At each sampling time Seedling and Graft plants were pruned to increase vigor. Three replicates were made at three weeks intervals for Graft and Seedling materials in 2018, and one replicate was made in 2019 including Tree material (twelve leaves were sampling). Each leaf was then photographed under binocular magnifier (Olympus SZ61, Schott KL 1500 LED, Olympus DP20). Pictures were further analyzed with the ImageJ® software (Schneider, Rasband, and Eliceiri 2012). Pictures were transformed in 8-bit grayscale and light intensity was measured on 5 areas of 0.03cm² on each leaf. Intensity differences between samples were evaluated using the R language by Kruskal-Wallis test. We first compare biological replicates from 2018 and from 2019 (Seed and Graft). Because there were no differences between biological replicates of Seedling from 2018 and 2019 and similarly to Graft sample from 2018 and 2019, we decide to only present result of the 2019 year which include Tree sample.

### DNA and RNA extraction

The youngest and completely opened leaf was sampled for each replicate. Sampling was performed as described in Table S4. The DNA was extracted using NucleoSpin Plant II kit (Macherey-Nagel, Hoerdt, France). The manufacturer’s recommendations were applied with the next modifications: at step 2a PL1 buffer quantity was raised to 800µL and PVP40 was added at 3% of final volume, suspension was then incubated 30min at 65°C under agitation. The lysate solution was centrifuged 2min at 11000g before transferring the supernatant in step 3. At step 4 PC buffer was raised to 900µL. In step 6 the first wash was decreased to 600µL and the third wash was raised to 300µL. An extra-centrifuge step was added after washing to remove ethanol waste from the column. In step 7 DNA was eluted twice in 55µL in total. The RNA was extracted using the NucleoSpin® RNA kit (Macherey-Nagel, Hoerdt, France) according the manufacturer’s protocol.

### Bisulfite sequencing and DMRs calling

Extracted DNA was precipitated in pure ethanol (70%), water (24%) and NaAc 3M (3%). After precipitation DNA was sent to Beijing Genomics Institute (Shenzhen, Guangdong 518083, China) in pure ethanol for whole genome bisulfite sequencing. DNA methylation data can be accessed on the Gene Expression Omnibus website under accession codes GSE138377. Bisulfite sequencing reads were mapped on GDDH13_V1.1 reference genome with Bsmap tool (Xi and Li 2009) to obtain methylation calling file. Methylation averages between samples were compared by student test using R (R Core Team 2016).

We called differentially methylated regions (DMRs) using a hidden Markov model (HMM)-based (Hagmann et al. 2015) approach as in Daccord *et al*., (2017). DMRs were calculated between Tree and Seedling samples and between Tree and Graft samples with the following parameters: coverage of 3, 200bp sliding windows with 100bp overlapping. DMRs files contain quality values such as p-value, average of standard deviation (SDA) and methylation differences. We empirically determined a threshold for each context using the DMR preview on a local JBrowse (Buels et al. 2016). This threshold was determined on SDA value (Supplemental tab. S5). Thresholds were determined in order to select the most reproducible DMRs within biological replicates (Supplemental tab. S6).

### Microarray

The *Malus domestica* array (Agilent-085275_IRHS_Malus_domestica_v1; GPL25795; Agilent, Foster City, CA, USA) was used for microarray analysis. Complementary DNA (cDNA) were synthesized and hybridized with the Low Input Quick Amp Labeling Kit, two-color (Agilent, Foster City, CA, USA). Two biological replicates were used. Each biological replicate represents one sample for Tree and Graft materials, and a pool of two samples for Seedling material. Hybridizations were performed on a NimbleGen Hybridization System 4 (mix mode B) at 42°C overnight. Slides were then washed, dried, and scanned at 2 µm resolution. NimbleGen MS 200 v1.2 software was used for microarray scans, and the Agilent Feature Extraction 11.5 software was used to extract pair-data files from the scanned images. We used the dye switch approach for statistical analysis as described in Depuydt *et al*., (2009). Analyses were performed using the R language (R Development Core Team, 2009); data were normalized with the lowess method, and differential transcription analyses were performed using the lmFit function and the Bayes moderated t test using the package LIMMA (Smyth, Michaud, and Scott 2005). Transcriptomic data are available in Gene Expression Omnibus website, with the accession GSE138491.

### RT-QPCR microarray validation

Extracted mRNA was treated by DNAse with the RQ1 RNase-Free DNase (Promega, Madison, WI, US) following the manufacturer’s protocol. The Moloney Murine Leukemia Virus Reverse Transcriptase was used to obtain cDNA from 1,2µg of total RNA, with oligot(dT) primers following the manufacturer’s protocol (Promega, Madison, WI, USA). For QPCR measurements, 2,5 µL of cDNA at the appropriate dilution were mixed in a final volume of 10µL with 5µL of quantitative PCR mastermix (MasterMix Plus for SYBR Green I with fluorescein; Eurogentec EGT GROUP, Seraing, Belgium), with 0.2µL of each primer (200nM final) and with 4,1µL of pure water. Primers were designed with Primer3Plus (Untergasser et al. 2007) and were used at their optimal concentration found thanks to reaction efficiency calculation (near to 100%) according to Pfaffl recommendations (Pfaffl 2001). Genes selected to validate the microarray data were selected in DEG lists in both comparisons (TvS and TvG) with 1) a high ratio value and 2) high intensities values. Accessions and primer sequences are indicated in figureS3A. Reaction was performed with a CFX connect Real time system (Bio-Rad, Hercules, CA, USA) using the following program: 95°C, 5 min; 35 cycles comprising 95°C for 3 s, 60°C for 45 s; 65°C, 5s and 90°C for 1 min, with real-time fluorescence monitoring. Melt curves were acquired at end of each run. Data were acquired and analyzed with CFX Maestro V1.1 (Bio-Rad, Hercules, CA, USA). Gene transcription levels were calculated using the 2^-ΔΔCt^ method and were corrected as recommended by Vandesompele *et al*., (2002) (Supplemental fig. S3B), with three reference genes: Actin (accession CV151413, MD14G1142600), Gapdh (accession CN494000, MD16G1111100), and Tubulin (accession CO065788, MD03G1004400) used for the calculation of a normalization factor.

### Differentially express transcript (DET) analysis

Differentially expressed transcripts were selected based on their p-value ≤ 1% (Supplemental tab. S7). For DET other than TE and miRNA a MapMan annotation (https://mapman.gabipd.org/home; version 3.5.0BETA), was performed, using GDDH13_1-1_mercator4 map file, in order to assign each DET to a BIN. DET not assigned to a BIN class were excluded. A representativeness percentage of each BIN class was then calculated in the comparisons TvS, TvG and in the intersection between the both comparisons. A MapMan enrichment analysis on the BIN class representativeness was performed and a BH correction was applied (Benjamini and Hochberg 1995) because of the high number of values. For DETEs, the TE classification (Daccord et al. 2017) was used in order to assign each DETE to a class.

### Association between DMR and DEG or DETE

DMRs and transcription level data (DEG or DETE) results were connected thanks to gene identifier. DMRs without associated DETs were removed. There is some redundancy of the gene or TE identification because many DMRs could be close. To avoid biases in our analysis we only kept DMR with the highest methylation variation to each gene or TE (Supplemental tab. S8).

## Accession numbers

GSE138492: global depository accession number comprising methylome and transcriptome data

GSE138377: bisulfite sequence data and methylation calling files

GSE138491: microarray data

## Large datasets

Supplemental table S6: DMRs list of TvS and TvG comparisons, including Gene- and TE-DMRs.

Supplemental table S7: DETs list of TvS and TvG comparison

Supplemental table S8: DEG- and DETE-DMRs list of TvS and TvG comparisons

## Acknowledgements

M. ORSEL-BALDWIN for GDDH13_1-1_mercator4 files processing. Region Pays de la Loire (FRANCE) to funding this work.

## Figure legends

**Supplemental figure S1: Methylation overview in GDDH13.** Density plot of DMRs on all GDDH13 genome in TvG. In red, DMRs in CG context, in blue CHG and in orange CHH. Each point represent number of DMRs in 50kb windows of genome.

**Supplemental figure S2: Jbrowse screenshoot** of two DEG-DMRs present in scatterplot (fig 7A) with DMRs in the promotor of genes highlight by a red dashed boxe.

**Supplemental figure S3:** (A) Q-PCR primers for micro array data validation. Indicated ratios came from micro array data in both comparisons. (B) Q-PCR validation of micro array data.

---

**Supplemental table S1:** Functional classes found in enrichment analysis on Mapman software. In red are indicate number of DEGs over transcribed in Tree sample, down regulated are in blue. “p-value” correspond to the p-value obtained in the enrichment analysis and corrected by the BH method.

| Category name | Bincode | Tree vs Seed |  |  | Tree vs Graft |  |  | Commons |  |  |  |
| --- | --- | --- | --- | --- | --- | --- | --- | --- | --- | --- | --- |
|  |  | Up | Down | p-value | Up | Down | p-value | Up | Down | other | p-value |
| Coenzyme metabolism | 7 | 0 | 19 | 2.56E-05 |  |  |  | 1 | 3 |  | 1.59E-02 |
| Secondary metabolism / terpenoids | 9.1 | 2 | 11 | 5.43E-03 |  |  |  |  |  |  |  |
| Secondary metabolism / phenolics / p-coumaroyl-coa synthesis | 9.2.1 |  |  |  | 0 | 2 | 7.50E-03 |  |  |  |  |
| Chromatin organization | 12 | 19 | 3 | 2.48E-05 | 1 | 14 | 6.03E-03 |  |  |  |  |
| Cell cycle | 13 |  |  |  | 5 | 39 | 9.05E-04 |  |  |  |  |
| Rna biosynthesis / transcriptional activation / SBP transcription factor | 15.7.18 | 8 | 0 | 3.37E-03 |  |  |  |  |  |  |  |
| Rna biosynthesis / organelle machineries | 15.9 |  |  |  |  |  |  | 1 | 5 |  | 1.50E-02 |
| Rna processing | 16 |  |  |  | 5 | 30 | 3.53E-03 |  |  |  |  |
| Protein biosynthesis | 17 | 9 | 40 | 3.17E-03 | 5 | 29 | 2.90E-04 | 5 | 53 |  | 4.07E-12 |
| Protein modification / phosphorylation / TKL kinase superfamily | 18.8.1 |  |  |  |  |  |  | 1 | 9 | 1 | 6.55E-03 |
| Protein degradation / peptide tagging | 19.4 | 18 | 15 | 7.94E-03 | 16 | 7 | 5.10E-04 |  |  |  |  |
| Protein degradation / peptidase families / serine-type peptidase activities | 19.5.2 |  |  |  |  |  |  | 1 | 9 | 1 | 8.33E-04 |
| Cytoskeleton | 20 |  |  |  | 1 | 20 | 1.04E-06 |  |  |  |  |
| Cell wall | 21 |  |  |  | 13 | 0 | 1.58E-05 |  |  |  |  |
| Protein translocation / chloroplast / thylakoid membrane SRP insertion system | 23.1.7 |  |  |  |  |  |  | 1 | 7 | 1 | 1.50E-02 |
| Solute transport | 24 |  |  |  | 16 | 1 | 3.12E-06 |  |  |  |  |
| Enzyme classification | 50 | 30 | 84 | 4.24E-03 | 31 | 25 | 7.32E-03 |  |  |  |  |
| Not assigned | 35 |  |  |  | 235 | 617 | 7.50E-03 | 122 | 252 | 14 | 1.17E-02 |

**Supplemental table S2:** Count of DEG-DMRs in TvS and TvG comparisons. Here we included the unclassified gene class “35” not present in Fig. 6.

| Context | localization | Tree vs Seed |  |  | Tree vs Graft |  |  |
| --- | --- | --- | --- | --- | --- | --- | --- |
|  |  | Hypermethylated | Hypomethylated | Σ | Hypermethylated | Hypomethylated | Σ |
| CHH | head | 1394 | 55 | 1449 | 406 | 261 | 667 |
|  | body | 295 | 21 | 316 | 137 | 70 | 207 |
|  | tail | 684 | 31 | 715 | 228 | 137 | 365 |
| CHG | head | 0 | 2 | 2 | 0 | 3 | 3 |
|  | body | 1 | 5 | 6 | 0 | 2 | 2 |
|  | tail | 0 | 4 | 4 | 0 | 3 | 3 |
| CG | head | 1 | 5 | 6 | 0 | 7 | 7 |
|  | body | 3 | 4 | 7 | 0 | 4 | 4 |
|  | tail | 1 | 4 | 5 | 0 | 10 | 10 |
| | $\Sigma$ | 2379 | 131 | 2510 | 771 | 497 | 1268 |

**Supplemental table S3:** Count of DETE-DMRs in CHH context in TvS and TvG comparisons.

| Context | localization | Tree vs Seed |  |  | Tree vs Graft |  |  |
| --- | --- | --- | --- | --- | --- | --- | --- |
| | | Hypermethylated | Hypomethylated | $\Sigma$ | Hypermethylated | Hypomethylated | $\Sigma$ |
| CHH | head | 0 | 0 | 0 | 0 | 0 | 0 |
|  | body | 115 | 22 | 137 | 153 | 113 | 266 |
|  | tail | 10 | 1 | 11 | 5 | 6 | 11 |
| CHG | head | 0 | 0 | 0 | 0 | 0 | 0 |
|  | body | 0 | 3 | 3 | 1 | 28 | 29 |
|  | tail | 0 | 0 | 0 | 1 | 0 | 1 |
| CG | head | 0 | 0 | 0 | 0 | 0 | 0 |
|  | body | 1 | 6 | 7 | 1 | 22 | 23 |
|  | Tail | 0 | 0 | 0 | 0 | 1 | 1 |
| | $\Sigma$ | 126 | 32 | 158 | 161 | 170 | 331 |

**Supplemental table S4:** Resume of defined samples and details of sampling.

| Sample | Way of multiplication | Years of obtention | Years sampling | Numbers sample | Pooled | Numbers leaves per sample | Numbers leaves sampled per tree | Numbers sampled trees |
| --- | --- | --- | --- | --- | --- | --- | --- | --- |
| **Tree** | Grafting (Asexual) | 2005 | 2016 | 3 | Yes | 4 | 4 | 1 |
| **Seed** | Seedling (Sexual) | 2016 | 2016 | 4 | No | 1 | 1 | 4 |
| **Graft** | Grafting (Asexual) | 2016 | 2016 | 2 | Yes | 10 | 1 | 20 |

**Supplemental table S5:** Fixed threshold to filter DMRs calculated between each comparison. Threshold were empirically fixed by observation of methylation calling file in the Jbrowse software.

| DMRs | Standard deviation average threshold |  |  | p-value |
| --- | --- | --- | --- | --- |
|  | Tree | Seed | graft |  |
| CG -CHG | 0,07 | 0,11 | 0,07 | 1% |
| CHH | 0,05 | 0,05 | 0,05 |  |

